# Succession of *Bifidobacterium longum* strains in response to the changing early-life nutritional environment reveals specific adaptations to distinct dietary substrates

**DOI:** 10.1101/2020.02.20.957555

**Authors:** Magdalena Kujawska, Sabina Leanti La Rosa, Phillip B. Pope, Lesley Hoyles, Anne L. McCartney, Lindsay J Hall

## Abstract

Diet-microbe interactions play a crucial role in infant development and modulation of the early-life microbiota. The genus *Bifidobacterium* dominates the breast-fed infant gut, with strains of *B. longum* subsp. *longum* (*B. longum*) and *B. longum* subsp. *infantis* (*B. infantis*) particularly prevalent. Although transition from milk to a more diversified diet later in infancy initiates a shift to a more complex microbiome, specific strains of *B. longum* may persist in individual hosts for prolonged periods of time. Here, we sought to investigate the adaptation of *B. longum* to the changing infant diet. Genomic characterisation of 75 strains isolated from nine either exclusively breast- or formula-fed (pre-weaning) infants in their first 18 months revealed subspecies- and strain-specific intra-individual genomic diversity with respect to glycosyl hydrolase families and enzymes, which corresponded to different dietary stages. Complementary phenotypic growth studies indicated strain-specific differences in human milk oligosaccharide and plant carbohydrate utilisation profiles of isolates between and within individual infants, while proteomic profiling identified active polysaccharide utilisation loci involved in metabolism of selected carbohydrates. Our results indicate a strong link between infant diet and *B. longum* subspecies/strain genomic and carbohydrate utilisation diversity, which aligns with a changing nutritional environment: i.e. moving from breast milk to a solid food diet. These data provide additional insights into possible mechanisms responsible for the competitive advantage of this *Bifidobacterium* species and its long-term persistence in a single host and may contribute to rational development of new dietary therapies for this important developmental window.

## Introduction

Microbial colonisation shortly after birth is the first step in establishment of the mutualistic relationship between the host and its microbiota (*1-3*). The microbiota plays a central role in infant development by modulating immune responses, providing resistance to pathogens, and also digesting the early-life diet (*4-10*). Indeed, diet-microbe interactions are proposed to play a crucial role during infancy and exert health effects that extend to later life stages (*11-16*). The gastrointestinal tract of vaginally delivered full-term healthy infants harbours a relatively simple microbiota characterised by the dominance of the genus *Bifidobacterium* (*17*).

Breast milk is considered the gold nutritional standard for infants, which also acts as an important dietary supplement for early-life microbial communities, including *Bifidobacterium*. The strong diet-microbe association has further been supported by reports of differences in microbial composition between breast- and formula-fed infants (e.g. high versus low *Bifidobacterium* abundance) and related differential health outcomes between the two groups: e.g. increased instances of asthma, allergy and obesity in formula-fed infants (*18-24*).

The high abundance of *Bifidobacterium* in breast-fed infants has been linked to the presence of specific carbohydrate utilisation genes and polysaccharide utilisation loci (PULs) in their genomes, particularly the ones involved in the degradation of breast milk-associated human milk oligosaccharides (HMOs) (*8*). The presence of these genes is often species- and indeed strain-specific, and has been described in *B. breve, B. bifidum, B. longum, B. infantis*, and more rarely in *B. pseudocatenulatum (8, 25-27)*. However, previous studies have indicated co-existence of *Bifidobacterium* species and strains in individual hosts, resulting in interaction and metabolic co-operation within a single (HMO-associated) ecosystem (*1, 28*).

Transition from breastfeeding to a more diversified diet and the introduction of solid foods has been considered to initiate the development of a functionally more complex adult-like microbiome with genes responsible for degradation of plant-derived complex carbohydrates, starches, and xenobiotics, as well as production of vitamins (*29, 30*). Non-digestible complex carbohydrates such as inulin-type fructans (ITF), arabino-xylans (AX) or arabinoxylo-oligosaccharides (AXOS) in complementary foods have been proposed to potentially exert beneficial health effects through their bifidogenic and prebiotic properties and resulting modulation of the intestinal microbiota and metabolic end-products (*31-34*).

Despite the shift in microbiota composition during weaning, specific strains of *Bifidobacterium*, and *B. longum* in particular, have previously been shown to persist in individuals over time (*35, 36*). *B. longum* is currently recognised as four subspecies: *longum* and *infantis* (characteristic of the human gut microbiota), and *suis* and *suillum* (from animal hosts) (*37, 38*). It is considered the most common and prevalent species found in the human gut, with *B. longum* subsp. *infantis* detected in infants, and *B. longum* subsp. *longum* widely distributed in both infants and adults (*39, 40*). The differences in prevalence between the two subspecies, and the ability of infant, adult and elderly host to acquire new *B. longum* strains during a lifetime have been attributed to distinct bacterial carbohydrate utilisation capabilities and the overall composition of the resident microbiota (*41, 42*). However, longitudinal assessments of this species in single hosts over the course of changing dietary patterns are limited, and therefore further detailed studies are required.

Here, we investigate the adaptations of *Bifidobacterium* to the changing infant diet and examine a unique collection of *B. longum* strains isolated from nine infants across their first 18 months. We probed the genomic and phenotypic similarities between 62 *B. longum* strains and 13 *B. infantis* strains isolated from either exclusively breast-fed or formula-fed infants (pre-weaning). Our results indicate a strong link between host diet and *Bifidobacterium* species/strains, which appears to correspond to the changing nutritional environment. Genome flexibility of *B. longum* and nutrient preferences of specific strains may aid their establishment within individual infant hosts, and their ability to persist through significant dietary changes. These dietary changes (moving from breast milk to solid food) may also encourage acquisition of new *B. longum* sp. strains with different nutritional preferences. Overall, our findings provide important additional insights into mechanisms responsible for adaptation to a changing nutritional environment and long-term persistence within the early life gut.

## Results

Previous investigations into *B. longum* across the human lifespan have determined a broad distribution of this species, including prolonged periods of colonisation (*35, 36*). To gain insight into potential mechanisms facilitating these properties during the early-life window, we investigated the genotypic and phenotypic characteristics of *B. longum* strains within individual infant hosts in relation to diet (i.e. breast milk vs formula) and dietary stages (i.e. pre-weaning, weaning and post-weaning), following up on a longitudinal study of the infant faecal microbiota (*43, 44*). Faecal samples from exclusively breast-fed infants and exclusively formula-fed infants were collected regularly from 1 month to 18 months of age (*43*). The number of samples obtained from the breast-fed infants during the pre-weaning period was higher than that obtained from the formula-fed group, which may correlate with differences in weaning age (∼20.6 vs. ∼17 weeks old). Bacterial isolation was carried out on faecal samples, and the isolated colonies identified using ribosomal intergenic spacer analysis (*44*). Based on these results, 88 isolates identified as *Bifidobacterium* were selected for this study, 46 from five exclusively breast-fed infants (BF1-BF5, including identical twins BF3 and BF4) and 42 from four exclusively formula-fed infants (FF1-FF3 and FF5). Following sequencing and ANI analysis (**Supplementary Tables S1 & S2**), 75 strains were identified as *B. longum* sp. and included in further analysis, with 62 strains identified as *B. longum* subsp. *longum* (*B. longum*) and 13 strains identified as *B. longum* subsp. *infantis* (*B. infantis*) (**Figure 1a**).

**Figure 1.**
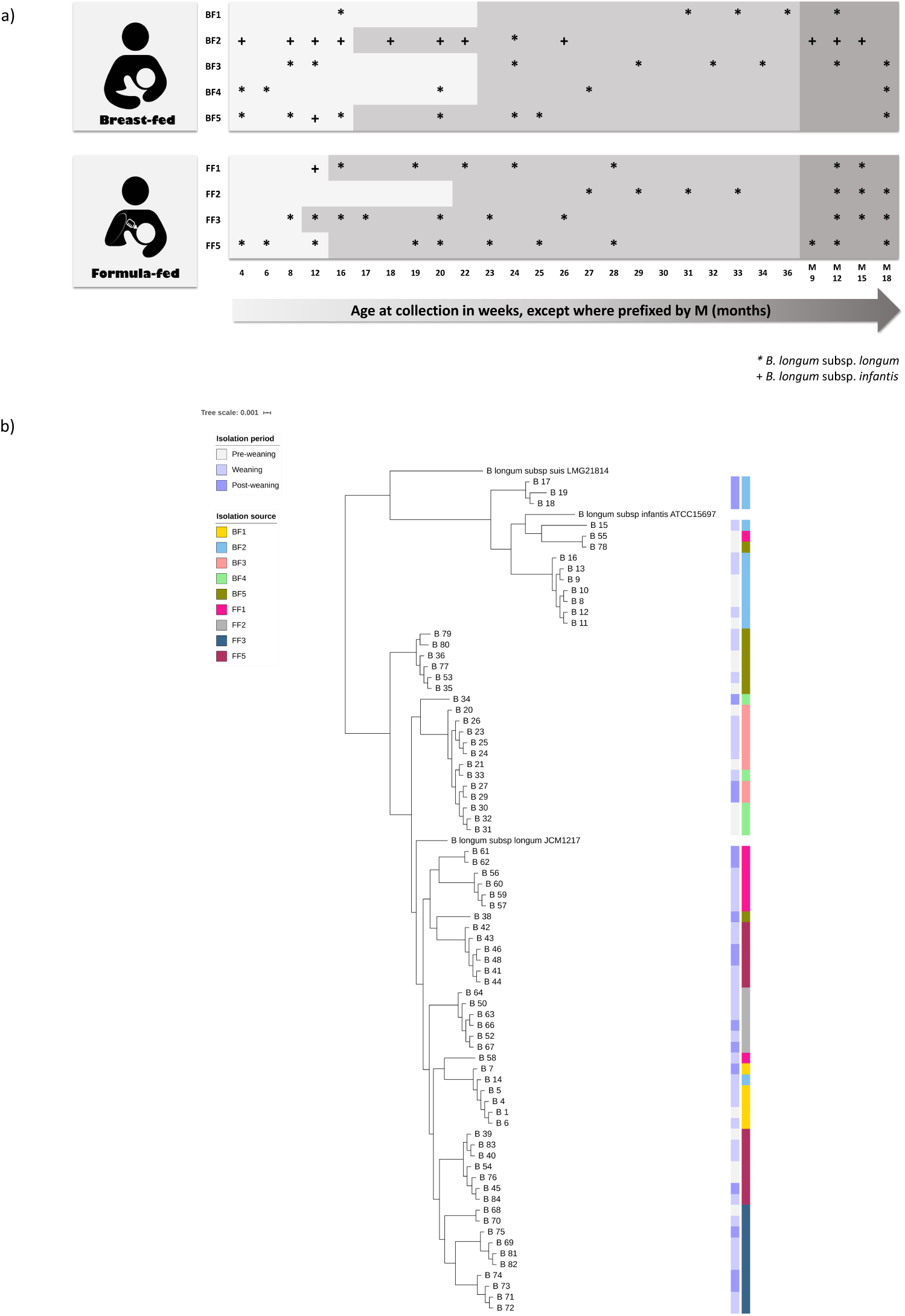
Identification and relatedness of *B. longum* strains. a) Sampling scheme and strain identification within individual breast-fed (BF1-BF5) and formula-fed (FF1-FF3 and FF5) infants based on average nucleotide identity values (ANI). b) Relatedness of *B. longum* strains based on core proteins. Coloured strips represent isolation period (pre-weaning, weaning and post-weaning) and isolation source (individual infants), respectively.

### General features of *B. longum* genomes

To determine possible genotypic factors facilitating establishment and persistence of *B. longum* in the changing early-life environment, we assessed the genome diversity of our strains. Sequencing generated between 12 and 193 contigs for each *B. longum* strain, with 98.6% of draft genomes (n=74) containing fewer than 70 contigs and one draft genome containing 193 contigs, yielding a mean of 66.95-fold coverage for strains sequenced on HiSeq (minimum 46-fold, maximum 77-fold) and 231-fold for the strain sequenced on MiSeq (**Supplementary Table S1**). The predicted genome size for strains identified as *B. longum* ranged from 2.21 Mb to 2.58 Mb, possessing an average G+C% content of 60.11%, an average predicted ORF number of 2,023 and number of tRNA genes ranging from 55-88. For strains identified as *B. infantis*, the predicted genome size ranged from 2.51 Mb to 2.75 Mb, with an average G+C% content of 59.69%, an average predicted ORF number of 2,280 and the number of tRNA genes ranging from 57 to 62.

### Comparative genomics

To identify *B. longum* strains among the sequenced isolates and assess the nucleotide-level genomic differences between isolates, we subjected their genomes to ANI analysis. Results (**Supplementary Table S2**) indicated that *B. longum* strains isolated from individual infant hosts displayed higher levels of sequence identity than strains isolated from different hosts. More specifically, pairwise identity values for strains isolated from infant BF3 showed the narrowest range (average value of 99.99±3.15e-5%), followed by infant FF2 strains (99.98±1.12e-4%), with infant BF2 strains having the broadest identity value range (averaging 99.13±7.8e-3%).

Next, we examined genetic diversity of newly sequenced *B. longum* strains and their relatedness to each other, and *B. longum* type strains, namely *B. longum* subsp. *longum* JCM 1217^T^, *B. longum* subsp. *infantis* ATCC 15697^T^ and *B. longum* subsp. *suis* LMG 21814^T^, based on the generated pangenome data. This analysis identified a total of 1002 genes as core genes present in at least 99% of the analysed *B. longum* subspecies genomes and allowed a clear distinction between *B. longum* subspecies (i.e. *longum* vs. *infantis*) based on the presence/absence of specific genes (**Supplementary Table S3**). Phylogenetic analysis performed on the *B. longum* core genome revealed that *B. longum* strains within each subspecies clustered mainly according to isolation source, i.e. individual infants, rather than dietary stage (i.e. pre-weaning, weaning and post-weaning) (**Figure 1b**). Interestingly, strains isolated from formula-fed baby FF5 clustered into two separate clusters, irrespective of the isolation period, suggesting presence of two highly related *B. longum* groups within this infant. Furthermore, strains isolated from identical twins BF3 and BF4 clustered together, indicating their close relatedness.

We next sought to identify whether specific components of the *B. longum* subspecies pangenome were enriched in infant hosts. Each candidate gene in the accessory genome was sequentially scored according to its apparent correlation to host diet (breast vs. formula) or dietary stage. A gene annotated as alpha-L-arabinofuranosidase, along with four other genes coding for hypothetical proteins, were predicted to be enriched in *B. longum* strains isolated from breast-fed infants. Alpha-L-arabinofuranosidases are enzymes involved in hydrolysis of terminal non-reducing alpha-L-arabinofuranoside residues in alpha-L-arabinosides and act on such carbohydrates as (arabino)xylans (*45, 46*). In addition, two genes coding for hypothetical proteins and a gene coding for Mobility protein A were overrepresented in strains isolated from formula-fed infants. We did not find any associations between specific genes and diet in *B. infantis*. Furthermore, no associations between genes and dietary stages were found in either *B. longum* or *B. infantis* (**Supplementary Table S4**).

As our strains were isolated from individual infants at different time points, we next sought to determine their intra-strain diversity; for this we used the first *B. longum* isolate from each infant as the ‘reference’ strain to which all other strains from the same infant were compared (**Figure 2**). Infants BF1, BF3 and FF2 had the lowest strain diversity; with respective mean pairwise SNP distances of 18.7±20.3 SNPs (mean±sd), 10.3±5.0 SNPs and 13.3±5.3 SNPs. These results suggest strains isolated from these infants may be clonal, indicating long-term persistence of *B. longum* within individual infant hosts despite early-life dietary changes. Surprisingly, analysis of strains isolated from breast-fed identical twins BF3 and BF4 revealed higher strain diversity in baby BF4 (mean pairwise SNP distance of 1034.5±1327.1 SNPs), compared to the highly similar strains in infant BF3 (i.e. 10.3±5.0 SNPs). Based on these results, we conducted SNP analysis on *B. longum* strains isolated from both babies and found that out of 13 strains analysed (n=8 from BF3 and n=5 from BF4), 12 isolated during pre-weaning, weaning and post-weaning appeared to be clonal (with mean pairwise SNP distance of 10.0±5.5 SNPs) and one strain from baby BF4 isolated post-weaning was more distant, with mean SNP distance of 2595.4±2.8 SNPs. The difference in strain diversity may relate to the fact that infant BF4 received a course of antibiotics during pre-weaning (at 14 weeks). *Bifidobacterium* counts were not detectable nor was any *Bifidobacterium*-specific PCR product for DGGE obtained from this infant during the antibiotic administration; however, both were obtained for the sample collected one week after antibiotic treatment completed (*44*). Furthermore, the presence of clonal strains in both babies suggests vertical transmission of *B. longum* from mother to both infants, or potential horizontal transmission between babies, consistent with previous reports (*42, 47-49*). *B. infantis* strains isolated from infant BF2 showed the highest strain diversity, with the mean pairwise SNP distance of 9030.9±8036.6 SNPs. Seven strains isolated during both pre-weaning and weaning periods appeared to be clonal, with mean pairwise SNP distance of 6.3±1.6 SNPs, while four strains isolated during weaning and post-weaning were more distant, with mean pairwise SNP distance of 14983.5±4658.3 SNPs (**Supplementary Table S5**).

**Figure 2.**
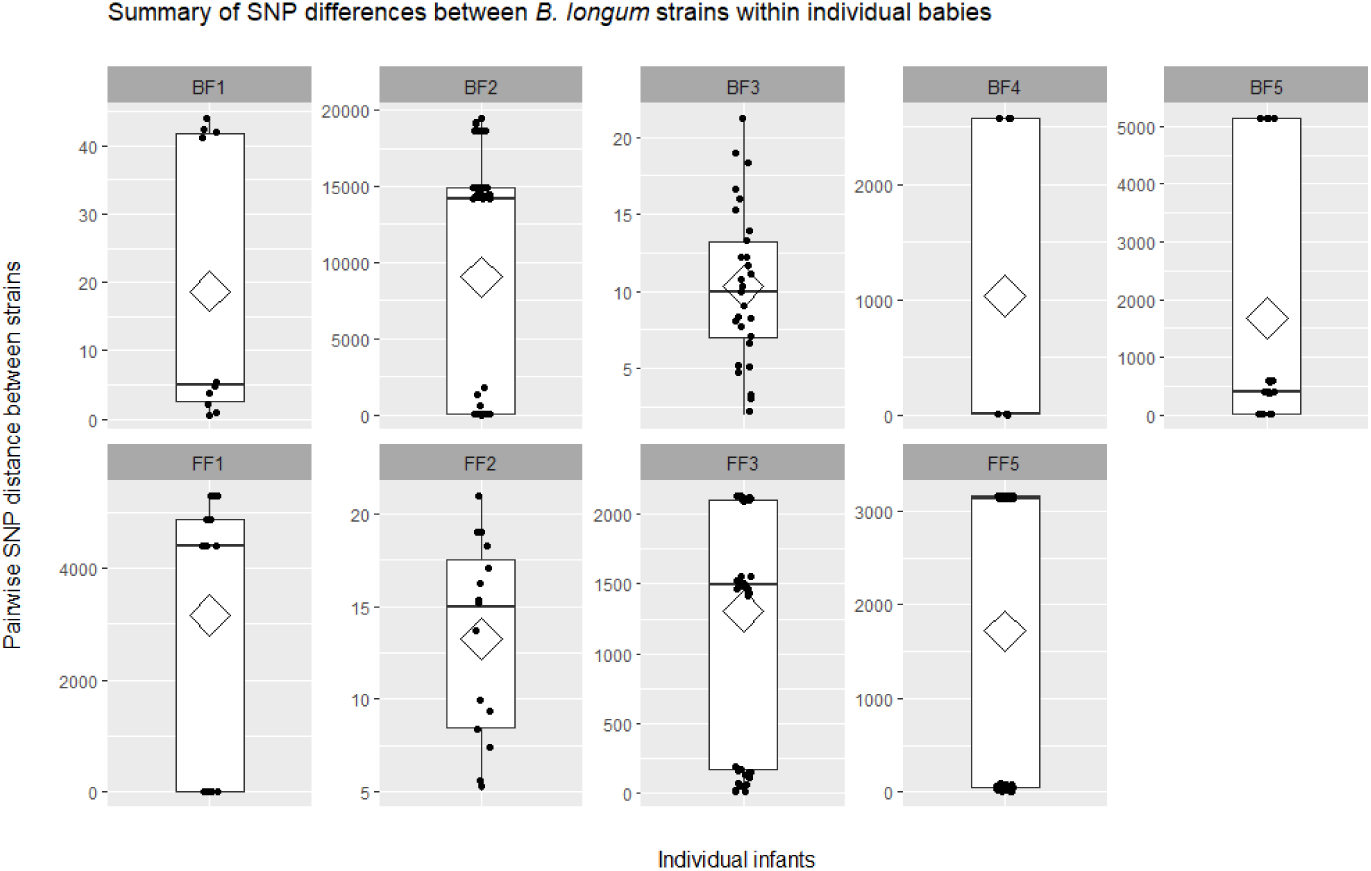
Pairwise SNP distances between *B. longum* strains of the same subspecies within individual infants. Individual points show data distribution, diamonds indicate the group mean, box plots show group median and interquartile range.

### Functional annotation of *B longum* subspecies genomes – carbohydrate utilisation

To assess genomic differences between our strains at a functional level, we next assigned functional categories to ORFs of each *B. longum* genome. Carbohydrate transport and metabolism was identified as the second most abundant category (after unknown function), reflecting the saccharolytic lifestyle of *Bifidobacterium* (**Supplementary Figure 1**) (*28, 50*). *B. longum* had a slightly higher proportion of carbohydrate metabolism and transport genes (11.39±0.31%) compared to *B. infantis* (10.20±0.60%), which is consistent with previous reports (*51, 52*). *B. longum* strains isolated during pre-weaning had a similar proportion of carbohydrate metabolism genes in comparison with the strains isolated post-weaning: 11.28±0.23% and 11.48±0.38%, respectively. Furthermore, we obtained similar results for *B. longum* strains isolated from breast- and formula-fed infants, with respective values of 11.41±0.21% and 11.38±0.38%. In contrast, *B. infantis* strains isolated pre-weaning had a lower proportion of carbohydrate metabolism genes in their genomes compared to the ones isolated post-weaning: 9.90±0.24% and 11.20±0.01%, respectively (**Supplementary Table S6**). These findings may indicate evolutionary adaptation of *B. infantis* strains to the changes in infant diet at early-life stages.

One of the major classes of carbohydrate-active enzymes comprises glycosyl hydrolases (GH), which facilitate glycan metabolism in the gastrointestinal tract (*53*). *Bifidobacterium* have been shown to possess an extensive repertoire of these enzymes, which allow them to adapt to the host environment through degradation of complex dietary and host-derived carbohydrates (*50*). We thus sought to investigate and compare the arsenal of GHs in *B. longum* sp. using dbCAN2. We identified a total of 36 different GH families in all *Bifidobacterium* strains. *B. longum* was predicted to contain 55 GH genes per genome on average (2.72 % of OFRs), while this number was lower for *B. infantis* strains - predicted to harbour an average of 37 GH genes per genome (1.62% of ORFs) (**Figure 3**).

**Figure 3.**
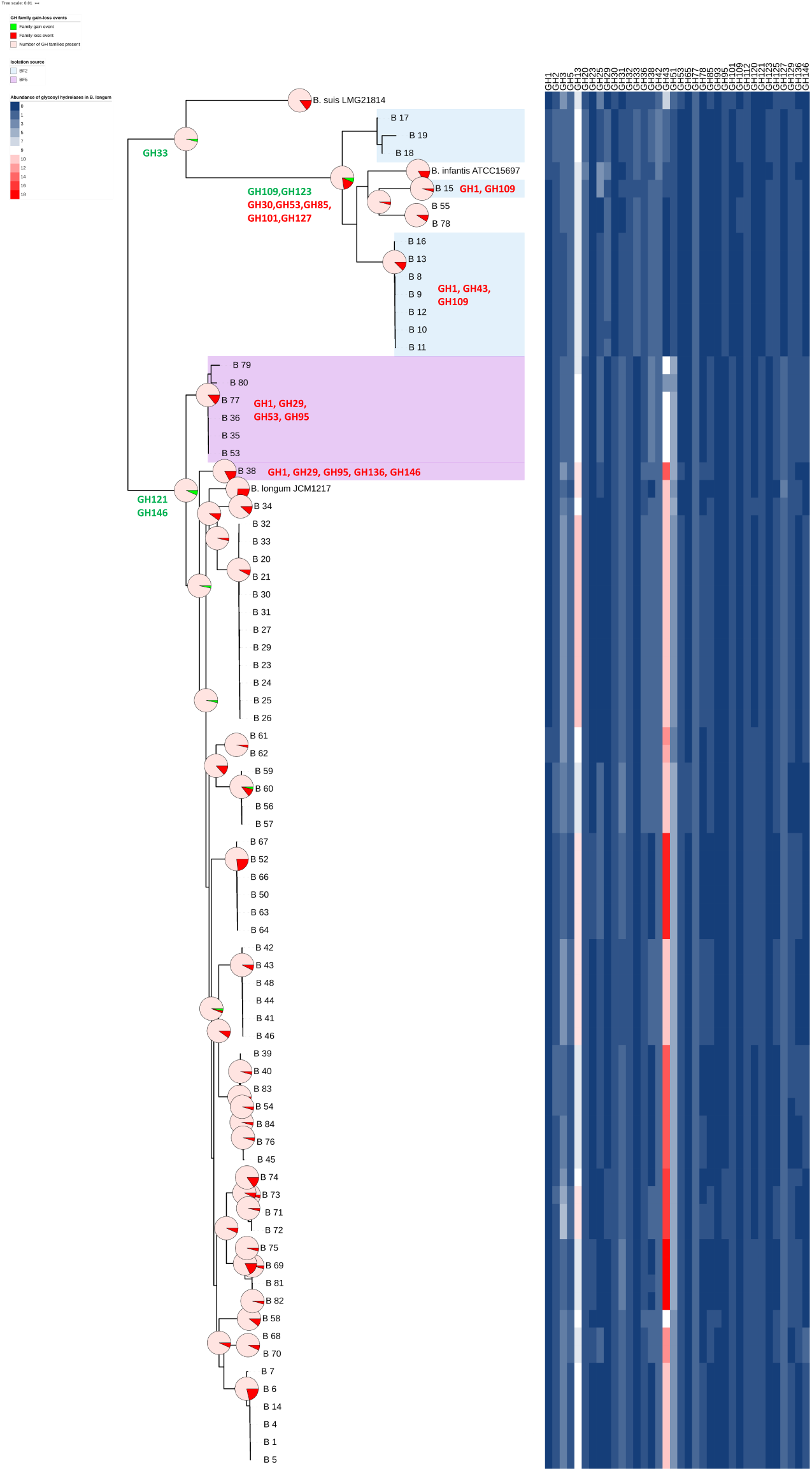
Gene-loss events and abundance of GH families within *B. longum* subspecies. Pie charts superimposed on the whole genome SNP tree represent predicted GH family gain-loss events within *B. longum* and *B. infantis* lineages. Due to the size of the tree, examples of detailed gain loss events have been provided for main lineages, as well as baby BF2 (strains highlighted with light blue) and BF5 (strains highlighted with light purple). Heatmap represents abundance of specific GH families predicted in analysed *B. longum* strains.

The predominant GH family in *B. longum* strains was GH43, whose members include enzymes involved in metabolism of complex plant carbohydrates such as (arabino)xylans (*54*), followed by GH13 (starch), GH51 (hemicelluloses) and GH3 (plant glycans) (*28, 55*).

Within the *B. longum* group, strains isolated during pre-weaning had a slightly lower mean number of GH genes compared to strains isolated post-weaning (54.46±2.81 vs. 56.85±2.77). Moreover, strains isolated from breast-fed babies contained an average of 53.96±3.82 GH genes per genome, while this number was slightly higher for strains isolated from formula-fed infants with 56.47±2.96 GH genes per genome. Further analysis revealed that differences in abundance of the predominant GH families in *B. longum* strains appeared to be intra-host-specific and diet-related. For example, strains isolated from breast-fed twins BF3 and BF4 pre-weaning had 11 GH43 genes per genome, while the pre-weaning strain from formula-fed baby FF3had 13 GH genes per genome predicted to belong to this GH family. Similarly, strains isolated from babies BF3 and BF4 post-weaning had 11 predicted GH genes, while the three strains isolated from infant FF3 were predicted to contain 16, 16 and 18 GH genes per genome, respectively (**Supplementary Table S7**).

In contrast, the most abundant GH family in *B. infantis* strains was GH13 (starch), followed by GH42, GH20 and GH38 (**Supplementary Table S7**). The GH42 family contains beta-galactosidases whose enzymatic activity ranges from lactose present in breast milk to galacto-oligosaccharides and galactans found in plant cell walls (*56, 57*). Members of GH20 family show hexosaminidase and lacto-*N*-biosidase activities, while family GH38 contains alpha-mannosidases (*28*). We also determined that, in contrast to *B. longum, B. infantis* strains harbour genes predicted to encode members of the GH33 family, which contains exo-sialidsaes (*28*). This finding suggested that *B. infantis* strains may have the ability to metabolise sialylated HMOs as well as utilise host mucins to release sialic acids and digest free sialic acid present in the gut.

Within the *B. infantis* group, strains isolated pre-weaning were predicted to contain an average of 34.83±0.4 GH genes per genome, while this number was higher for the strains isolated post-weaning (i.e. 43.00±0.00 GH genes per genome). Further analysis revealed that *B. infantis* strains isolated post-weaning contained families GH1 and GH43 that were absent in the strains isolated pre-weaning. The GH1 family contains enzymes such as beta-glucosidases, beta-galactosidases and beta-D-fucosidases active on a wide variety of (phosphorylated) disaccharides, oligosaccharides, and sugar–aromatic conjugates (*58*). In addition, the *B. infantis* strains isolated post-weaning harboured a higher number of genes predicted to belong to families GH42 and GH2 (enzymes active on a variety of carbohydrates) (*59*).

Members of the genus *Bifidobacterium* have previously been shown to contain GH genes involved in metabolism of various HMOs present in breast milk (*27, 60*). Alpha-L-fucosidases belonging to families GH29 and GH95 have been determined to show specificity towards fucosylated HMOs (*27, 61*), while lacto-*N*-biosidases and galacto-*N*-biose/lacto-*N*-biose phosphorylases members of GH20 and GH112 have been shown to be involved in degradation of isomeric lacto-*N*-tetraose (LNT) (*62*). We identified genes belonging to GH29 and GH95 in all our *B. infantis* strains, as well as four *B. longum* strains isolated from formula-fed baby FF3. Furthermore, we found GH20 and GH112 genes in all our *B. infantis* and *B. longum* strains (**Supplementary Table S7**).

Overall, these findings suggest differences in general carbohydrate utilisation profiles between *B. longum* and *B. infantis*. The presence of genes involved in utilisation of different carbohydrates, including HMOs in our strains, suggests the adaptation of *Bifidobacterium* to a changing early-life nutritional diet, which may be a factor facilitating establishment of these bacteria within individuals during infancy.

### Prediction of gain and loss of GH families in *B. longum*

Given the differences in the carbohydrate utilisation profiles between *B. longum* and *B. infantis*, we next investigated the acquisition and loss of GH families within the two subspecies. For this purpose, we additionally predicted the presence of GH families in type strains *B. longum* subsp. *longum* JCM 1217^T^, *B. longum* subsp. *infantis* ATCC 15697^T^ and *B. longum* subsp. *suis* LMG 21814^T^ with dbCAN2 and generated a whole genome SNP tree to reflect gene loss/gain events more accurately (**Figure 3, Supplementary Table S8**). Both *B. longum* and *B. infantis* lineages appear to have acquired GH families (when compared to the common ancestor of the phylogenetic group), with the *B. longum* lineage gaining two GH families (GH121 and GH146) and the *B. infantis* lineage one GH family (GH33). Within the *B. infantis* lineage, which also contains the *B. suis* type strain, the *B. infantis* taxon has further acquired two and lost five GH families. These findings suggest that the two human-related subspecies have followed different evolutionary paths, which is consistent with our observation of differences between *B. longum* and *B. infantis* resulting from phylogenomic analyses. Intriguingly, strain adaptation to the changing host environment (i.e. individual infant gut) seems to be driven by loss of specific GH families (**Figure 3**). For example, *B. infantis* strains isolated during pre-weaning and weaning from baby BF2 appear to be missing up to three GH families (GH1, GH43 and GH109) present in strains isolated post-weaning. Lack of family GH43 (containing enzymes involved in metabolism of a variety of complex carbohydrates, including plant-derived polysaccharides) in early-life *B. infantis* strains may explain nutritional preference of this subspecies for an HMO-rich diet. Similarly, we observed differential gene loss events in *B. longum* strains from individual hosts. For example, all strains isolated from baby BF5 appear to lack GH families GH1, GH29 and GH95. However, strains isolated pre-weaning additionally lacked GH53 family, which includes endogalactanases shown to be involved in liberating galactotriose from type I arabinogalactans in *B. longum* (*63*). In contrast, strain B_38 isolated from this infant (BF5) post-weaning appears to have lost families GH136 and GH146. Interestingly, members of family GH136 are lacto-*N*-biosidases responsible for liberating lacto-*N*-biose I from LNT, an abundant HMO unique to human milk (*64*). Overall, the presence of intra-individual and strain-specific GH family repertoires in *B. longum* suggests their adaptation to host-specific diet. The presence of strains with different GH content at different dietary stages further indicates potential acquisition of new *Bifidobacterium* strains with nutrient-specific adaptations in response to the changing infant diet.

### Phenotypic characterisation of carbohydrate utilisation

*Bifidobacterium longum* has previously been shown to metabolise a range of carbohydrates, including dietary and host-derived glycans (*65, 66*). Given the predicted differences in carbohydrate metabolism profiles between *B. longum* and *B. infantis*, and to understand strain-specific nutrient preferences of our strains, we next sought to determine their glycan fermentation capabilities. We performed growth assays on 49 representative strains from all nine infants, cultured in modified MRS supplemented with selected carbohydrates as the sole carbon source. For these experiments, we chose both plant- and host-derived glycans that we would expect to constitute components of the early-life infant diet (*67*). Although all *B. longum* strains were able to grow on simple carbohydrates (i.e. glucose and lactose), we also observed subspecies-specific complex carbohydrate preferences, consistent with bioinformatic predictions (**Figure 4**). To represent host-derived carbohydrates, we selected 2’-fucosyllactose (2’-FL) and lacto-*N*-neotetraose (LNnT) as examples of HMOs found in breast milk. Out of the tested isolates, all *B. infantis* strains were able to metabolise 2’-FL, as were three *B. longum* strains isolated from a formula-fed baby FF3 during weaning and post-weaning (**Figure 4**). These results supported the computational analysis and the identification of genes potentially involved in degradation of fucosylated carbohydrates in the genomes of these isolates (GH29 and GH95). Although bioinformatics identified the presence of genes involved in metabolism of isomeric LNT in all our strains (GH20 and GH112), LNnT metabolism in *B. infantis* was strain-specific, with most strains showing moderate to high growth rates. Out of *B. longum* strains, B_25 (isolated during weaning from breast-fed baby BF3) also showed robust growth on LNnT. Furthermore, this strain was the only strain out of the 49 tested that showed growth on cellobiose and, in contrast to all other *B. longum* strains, was not able to metabolise plant-derived arabinose and xylose despite the predicted presence of genes involved in metabolism of monosaccharides (GH43, GH31, GH2). Additionally, both *B. longum* and *B. infantis* strains showed varying degrees of growth performance on mannose, while none of the tested strains were able to grow on arabinogalactan, pectin or rhamnose (**Figure 4**).

**Figure 4.**
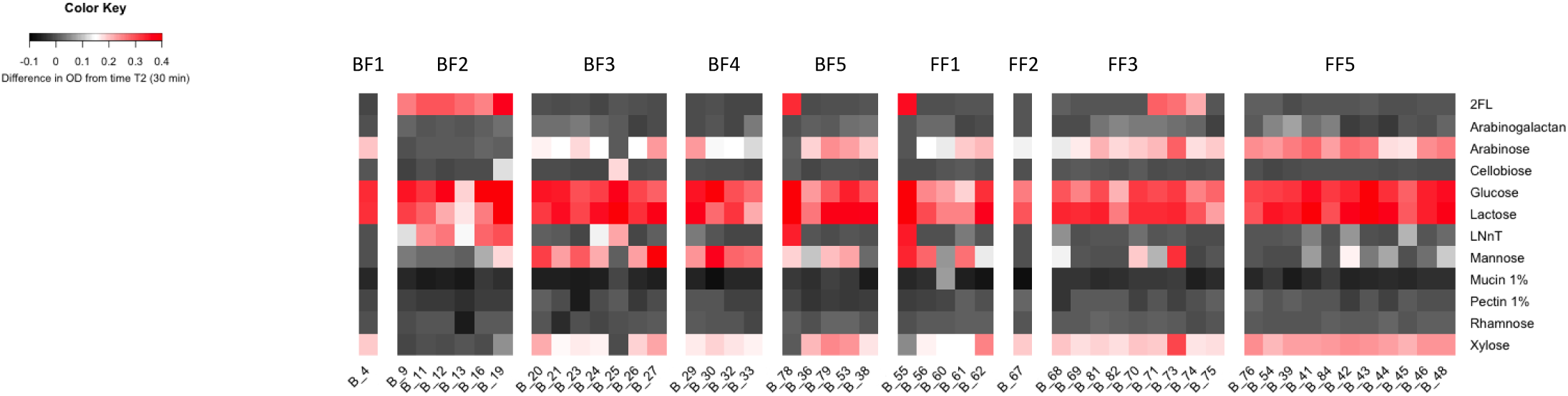
Growth performance of *B. longum* strains on different carbon sources. Heatmap displays the difference in average growth of triplicates between T_2_ (30 min) and T_end_ (48 hours).

To further characterise strains identified above for putative carbohydrate degradation genes, we performed carbohydrate uptake analysis and proteomics. *B. longum* strain B_25, from one of the breast-fed identical twins that showed growth on LNnT and cellobiose, and formula-fed strain B_71 which was able to grow on 2’-FL, were chosen. Supernatant from these cultures was initially subjected to high-performance anion-exchange chromatography (HPAEC) to evaluate the carbohydrate-depletion profiles (**Figure 5**). In all three cases, the chromatograms showed complete utilisation of the tested carbohydrates and absence of any respective degradation products in the stationary phase culture. The depletion of cellobiose by B_25 and 2’-FL by B_71 occurred in the early exponential phase while LNnT was still detected in the culture supernatant until the late exponential phase of growth, suggesting that cellobiose and 2’-FL were internalised more efficiently than LNnT. We next determined the proteome of B_25 and B_71 when growing on cellobiose, LNnT and 2’-FL compared to glucose (**Figure 5a-c & Supplementary Table S9**). The top 10 most abundant proteins in the cellobiose proteome of B_25 included three beta-glucosidases belonging to GH3 family, as well as a homologue of transport gene cluster previously shown to be upregulated in *B. animalis* subsp. *lactis* Bl-04 during growth on cellobiose (**Figure 5a & Supplementary Table S10**) (*68*). Among the three β-glucosidases, B_25_00240 showed 98% sequence identity to the structurally characterized BlBG3 from *B. longum*, which has been shown to be involved in metabolism of the natural glycosides saponins (*69*). B_25_01763 and B_25_00262 showed 46% identity to the β-glucosidase Bgl3B from *Thermotoga neapolitana* (*70*) and 83% identity to BaBgl3 from *B. adolescentis* ATCC 15703 (*71*), respectively, two enzymes previously shown to hydrolyse cello-oligosaccharides. With respect to LNnT metabolism by the same strain, the most abundant proteins were encoded by genes located in two PULs (B_25_00111-00117 and B_25_00130-00133) with functions compatible with LNnT import, degradation to monosaccharides and further metabolism. The PULs contain the components of an ABC-transporter (B_25_00111-00113), a predicted intracellular GH112 lacto-*N*-biose phosphorylase (B_25_00114), an *N*-acetylhexosamine 1-kinase (B_25_00115) and enzymes involved in the Leloir pathway. All these proteins were close homologues to proteins previously implicated in the degradation of LNT/LNnT by type strain *B. infantis* ATCC 15697^T^ (*72*) (**Figure 5b & Supplementary Table S10**). Interestingly, all clonal strains isolated from twin babies BF3 and BF4 also contained close homologues of all the above-mentioned genes in their genomes, in some cases identical to those determined in B_25; however, only strain B_25 was able to grow on cellobiose and LNnT. Growth of B_71 on 2’-FL corresponded to increased abundance of proteins encoded by the PUL B_71_00973-00983. These proteins showed close homology to proteins described for *B. longum* SC596 and included genes for import of fucosylated oligosaccharides, fucose metabolism and two α-fucosidases belonging to the families GH29 and GH95 (**Figure 5c & Supplementary Table S10**) (*27*).

**Figure 5.**
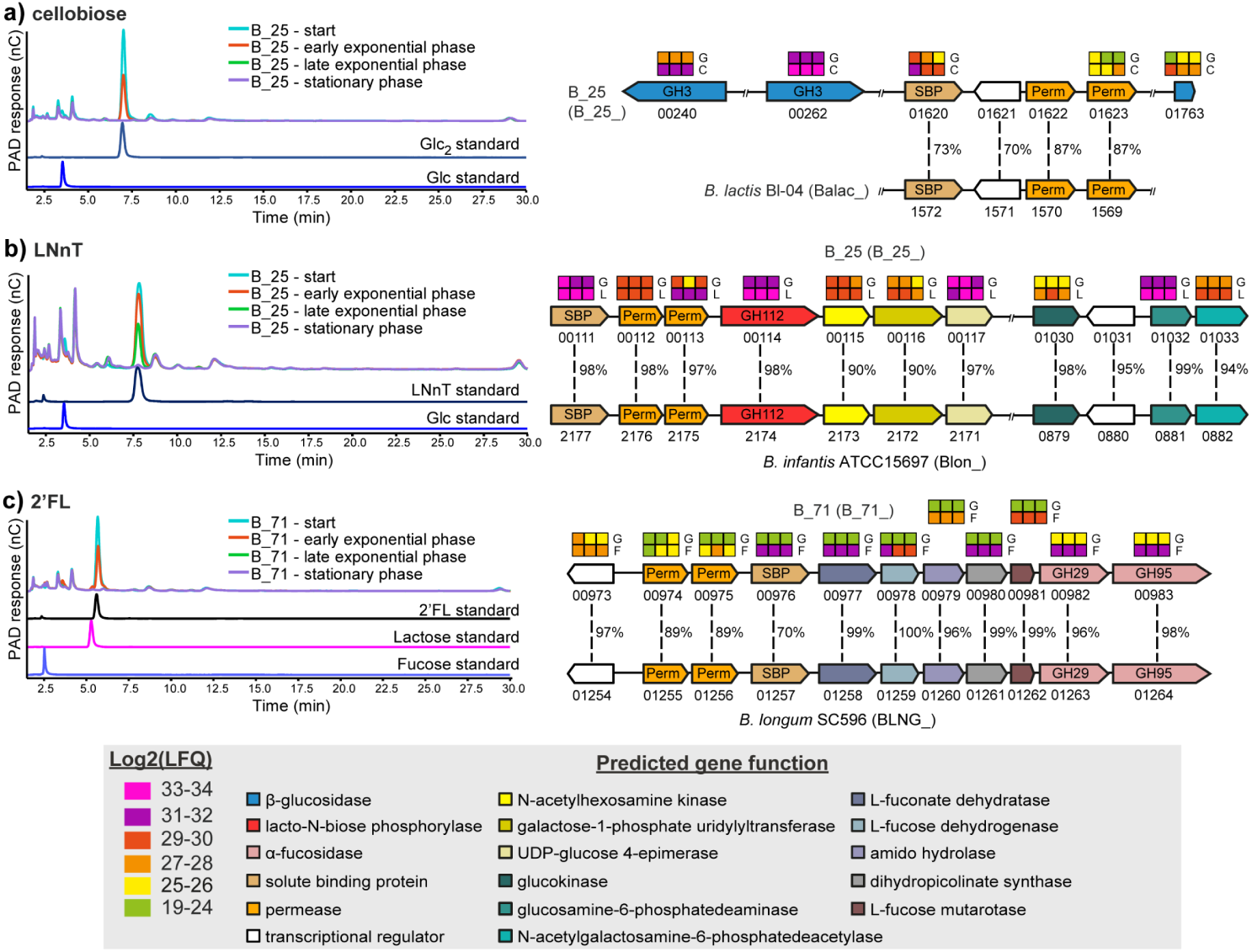
HPAEC-PAD traces showing mono-, di- and oligo-saccharides detected in the supernatant of either B_25 or B_71 single cultures during growth in mMRS supplemented with (a) cellobiose; (b) LNnT; (c) 2’-FL. The data are representative of biological triplicates. Abbreviations: LNnT, Lacto-*N*-neotetraose; Glc, glucose; Glc2, cellobiose; 2’-FL, 2′-fucosyllactose. Panel on the right shows (a) cellobiose; (b) LNnT; (c) 2’-FL utilization clusters in B_25 and B_71 and proteomic detection of the corresponding proteins during growth on HMOs. Heat maps above genes show the LFQ detection levels for the corresponding proteins in triplicates grown on glucose (G); cellobiose (C); LNnT (L); and 2’-FL (F). Numbers between genes indicate percent identity between corresponding genes in homologous PULs relative to strains B_25 and B_71. Numbers below each gene show the locus tag in the corresponding genome. Locus tag numbers are abbreviated with the last numbers after the second hyphen (for example B_25_XXXXX). The locus tag prefix for each strain is indicated in parenthesis beside the organism name.

## Discussion

High abundance of *Bifidobacterium* in early infancy is strongly linked to availability of nutrients, with dominance in breast-fed infants correlated with enrichment of genes required for the degradation of HMOs present in breast milk, while the transition to solid foods during weaning has been linked to genes involved in degradation of complex plant-derived carbohydrates (*3, 29, 64*). *Bifidobacterium longum* species appear to be widely distributed in individuals at different life stages, which may correlate with the abundance of genes responsible for carbohydrate metabolism (*35, 42*). In this study, we aimed to investigate the adaptations of *B. longum* to the changing infant diet during the early-life developmental window. We analysed the intra-subspecies genomic diversity of 75 *B. longum* strains isolated from nine individual infant hosts at different dietary stages, with focus on their potential carbohydrate metabolism capabilities, and determined their growth performance on different carbohydrates as sole carbon sources. Our results indicate intra-individual and diet-related differences in genomic content of analysed strains, which links to their ability to metabolise specific dietary components.

Our comparative genomic analysis indicates that clonal strains of *B. longum* sp. can persist in individuals through infancy, for at least 18 months, despite significant changes in diet during weaning, which is consistent with previous reports (*35, 42*). Concurrently, new strains (that display different genomic content and potential carbohydrate metabolism capabilities) can be acquired, possibly in response to the changing diet. Initial vertical acquisition of *Bifidobacterium* from mother to newborn babies has been well documented (*48, 49, 73*); however, details of strain transmission events in later life are currently unclear. Work of Odamaki et al. (*42*) identified person-to-person horizontal transmission of a particular *B. longum* strain between members of the same family, and suggested direct transfer, common dietary sources or environmental reservoirs, such as family homes (*74*), as potential vehicles and routes for strain transmission. Our results showed the presence of clonal strains in identical twins BF3 and BF4, which may have resulted from maternal transfer. However, potential strain transmission between these infants living in the same environment may also occur. Furthermore, wider studies involving both mothers and twin babies (and other siblings) could provide details on the extent, timing and location of transmission events between members of the same household.

Another aspect of comparative genomic analysis involved *in-silico* prediction of genes belonging to GH families. This analysis revealed genome flexibility within *B. longum* sp., with differences in GH family content between strains belonging to the same subspecies as described previously; *B. infantis* predominantly enriched in GH families implicated in the degradation of host-derived breast milk-associated dietary components like HMOs and *B. longum* containing GH families involved in the metabolism of plant-derived substrates (*28, 55*). Within the *B. infantis* group, we identified subspecies-specific differences in GH content between pre- and post-weaning strains, which indicates adaptation to the changing infant diet. Moreover, we observed differences in the number of genes belonging to the most abundant GH families (e.g. GH43) between breast-fed and formula-fed strains at different dietary stages, which can be linked to nutrient availability. Surprisingly, we computationally and phenotypically identified closely related weaning and post-weaning *B. longum* strains capable of metabolising HMOs (i.e. 2’-FL) in a formula-fed baby that only received standard non-supplemented (i.e. no prebiotics or synthetic HMOs) formula. However, these data should be carefully interpreted, since our collection only contains one bacterial strain per time point. In addition, analysis of strains belonging to *B. infantis* group was performed based on strains from only one breast-fed baby, which is a further caveat of the study.

Recorded phenotypic data support the results of genomic analyses and further highlight differences in carbohydrate utilisation profiles between and within *B. longum* and *B. infantis*. The ability of *Bifidobacterium*, especially *B. infantis*, to grow on different HMOs indicates their adaptation to an HMO-rich diet, which may be a factor facilitating their establishment within hosts at early-life stages. Similarly, *B. longum* preference for plant-based nutrients may be influencing their ability to persist within individual hosts through significant dietary changes. Differential growth of strains that are genotypically similar on various carbohydrate substrates and the ability of formula-fed strains to metabolise selected HMOs suggest that *Bifidobacterium* possess an overall very broad repertoire of genes for carbohydrate acquisition and metabolism that may be differentially switched on and off in response to the presence of specific dietary components (*75, 76*). Another explanation for these results may be a potential influence of the intra-individual environment on epigenetic mechanisms in these bacteria. One potential factor involved in this process may be a cooperative effort among *Bifidobacterium* in the early-life microbiota supported by cross-feeding activities between species and strains (*1, 28*).

Glycan uptake analysis and proteomic investigation allowed us to determine mechanisms which selected *B. longum* strains employ to metabolise different carbohydrates. A common feature, based on the predicted activity of the most abundant proteins detected during grown on the three substrates (cellobiose, LNnT and 2’-FL), was that they were all imported and “selfishly” degraded intracellularly, therefore limiting release of degradation products that could allow cross-feeding by other gut bacteria. This is in line with the carbohydrate uptake analysis, where no peak for cellobiose, LNnT and 2’-FL degradation products could be detected. Cellobiose uptake in B_25 occurs via a mechanism similar to *B. animalis* subsp. *lactis* Bl-04 (*68*); cellobiose hydrolysis appears to be mediated by the activity of three intracellular β-glucosidases, although further confirmatory biochemical characterization of these enzyme is still required. B_25 was observed to utilize LNnT using a pathway similar to that described in *B. longum* subsp. i*nfantis* whereby LNnT is internalized via an ABC-transporter (B_25_00111-00113) followed by intracellular degradation into constituent monosaccharides by a GH112 (B_25_00114) and an *N*-acetylhexosamine 1-kinase (B_25_00115). LNnT degradation products are further metabolized to fructose-6-phosphate by activities that include B_25_00116-00117 (galactose-1-phosphate urydyltranferase, UDP-glucose 4-epimerase, involved in the Leloir pathway) and B_25_01030-01033 (for metabolism of *N*-acetylgalactosamine) prior to entering the *Bifidobacterium* genus-specific fructose-6-phosphate phosphoketolase (F6PPK) pathway (*72*). B_71 is predicted to deploy an ABC-transporter (B_71_00974-00976) that allows uptake of intact 2’-FL that is subsequently hydrolysed to L-fucose and lactose by the two predicted intracellular α-fucosidases GH29 (B_71_00982) and GH95 (B_71_00983). L-fucose is further metabolized to L-lactate and pyruvate, via a pathway of non-phosphorylated intermediates that include activities of L-fucose mutarotase (B_71_00981), L-fucose dehydrogenase (B_71_00978), L-fuconate hydrolase (B_71_00977) as previously described for *B. longum* subsp. *longum* SC596 (*27*). Considering that the proteins encoded by the aforementioned genes are located in the cellobiose, LNnT and 2’-FL PULs that share high similarity and similar organization with those found in equivalent systems in other *B. longum* and *B. animalis*, it is reasonable to suggest that the PULs are related and may be the results of horizontal gene transfer events between *B. longum*/*B. animalis* members residing in the infant gut microbiota. Collectively, these data reflect inter- and intra-host phenotypic diversity of *B. longum* ssp. in terms of their carbohydrate degradation capabilities and suggest that intra-individual environment may influence epigenetic mechanisms in *Bifidobacterium*, resulting in differential growth on carbohydrate substrates.

In conclusion, this research provides new insight into distinct genomic and phenotypic abilities of *B. longum* species and strains isolated from the same individuals during the early-life developmental window by demonstrating that subspecies- and strain-specific differences between members of *B. longum* sp. in infant hosts can be correlated to their adaptation at specific age and diet stages.

## Materials and methods

### Bacterial isolates

Infants were recruited between 2005 and 2007: five were exclusively breast-fed and four were exclusively formula-fed. Faecal samples were obtained from infants at specific intervals during the first 18 months of life. For inclusion in the study, infants had to meet the following criteria: have been born at full-term (>37 weeks gestation); be of normal birth weight (>2.5 kg); be <5 weeks old and generally healthy; and be exclusively breast-fed or exclusively formula-fed [SMA Gold or SMA White (Wyeth Pharmaceuticals), to avoid supplemented formulae and to keep consistency within the formula group]. The mothers of the breast-fed infants had not consumed any antibiotics within the 3 months prior to the study and had not taken any prebiotics and/or probiotics. Ethical approval was obtained from the University of Reading Ethics Committee (*43*). *Bifidobacterium* strains (n=88) were isolated from healthy infants (**Supplementary Table S1**), either exclusively breast-fed (BF) or formula-fed (FF), and originally identified using ribosomal intergenic spacer analysis (*44*).

### DNA extraction, whole-genome sequencing, assembly and annotation

Phenol-chloroform method used for genomic DNA extraction as described previously (*1*). DNA isolated from pure bacterial cultures was subjected to multiplex Illumina library preparation protocol followed by sequencing on Illumina HiSeq 2500 platform (n=87) at the Wellcome Trust Sanger Institute (Hinxton, UK) or Illumina MiSeq (n=1) at Quadram Institute Bioscience (Norwich, UK) with read length of PE125 bp and PE300 bp, respectively, with an average sequencing coverage of 66.95-fold for isolates sequenced on HiSeq (minimum 46-fold, maximum 77-fold) and 231-fold for the isolate sequenced on MiSeq (**Supplementary Table S1**). Sequencing reads were checked for contamination using Kraken v1.1 (MiniKraken) (*77*) and pre-processed with fastp v0.20 (*78*) before assembling using SPAdes v3.11 with “careful” option (*79*). Contigs below 500bp were filtered out from the assemblies. Incorrectly assembled sequences were removed from further analysis (n=3). Additionally, publicly available assemblies of *Bifidobacterium* type strains (n=70) (**Supplementary Table S1)** were retrieved from NCBI Genome database and all genomes were annotated with Prokka v1.13 (*80*). The draft genomes of 75 *B. longum* isolates have been deposited to GOLD database at https://img.jgi.doe.gov, GOLD Study ID: Gs0145337.

### Phylogenetic analysis

Python3 module pyANI v0.2.7 with default BLASTN+ settings was employed to calculate the average nucleotide identity (ANI) (*81*). Species delineation cut-off was set at 95% identity (*82*) and based on that only sequences identified as *Bifidobacterium longum* subspecies were selected for further analysis (n=75) (**Supplementary Table S2**).

General feature format files of *B. longum* strains were inputted into the Roary pangenome pipeline v.3.12.0 to obtain core-genome data and the multiple sequence alignment (msa) of core genes (Mafft v7.313) (*83, 84*). SNP analysis of strains from individual infants was performed using Snippy v4.2.1 (*85*) and the resulting msa was passed to the recombination removal tool Gubbins(*86*). Alignments resulting from all previous steps were cleaned from poorly aligned positions using manual curation and Gblocks v0.9b where appropriate (*87*). The core-genome tree was generated using FastTree v2.1.9 using the GTR model with 1000 bootstrap iterations (*88*). Snp-dists v0.2 was used to generate pairwise SNP distance matrix between strains within individual infants (*89*). Altogether, the results of the SNP analysis reflected ANI results, showing that pairwise sequence identities were inversely proportional to pairwise SNP distances in *B. longum* subspecies isolates recovered from individual hosts.

### Functional annotation and genome-wide association study analysis

Scoary v1.6.16 with Benjamini Hochberg correction (*90*) was used to associate subsets of genes with specific traits – breast-fed, formula-fed, pre-weaning, weaning and post-weaning. The p-value cut-off was set to <1e-5, sensitivity cut-off to ≥70 % and specificity cut-off to ≥90 % to report the most overrepresented genes. Functional categories (COG categories) were assigned to genes using EggNOG-mapper v0.99.3, based on the EggNOG database (bacteria) (*91*) and the abundance of genes involved in carbohydrate metabolism was calculated. As most *B. infantis* strains (12 out of 13) were isolated from breast-fed infants, we did not compare abundances of carbohydrate metabolism genes in breast-fed and formula-fed groups for this subspecies. Standalone version of dbCAN2 (v2.0.1) was used for CAZyme annotation (*92*). Glycosyl hydrolase (GH) gain-loss events were predicted using Dollo parsimony implemented in Count v9.1106 (*93*). Snippy v4.2.1 with the “--ctgs” option, SNP-sites v2.3.3 (*94*) and FastTree v2.1.9 (GTR model with 1000 bootstrap iterations) were used to generate the whole genome SNP tree.

### Carbohydrate utilisation

To assess the carbohydrate utilisation profile, *Bifidobacterium* (1%, v/v) was grown in modified (m)MRS (pH 6.8) supplemented with cysteine HCl at 0.05% and 2% (w/v) of selected carbohydrates (HMOs obtained from Glycom, Hørsholm, Denmark) as described previously (*1*), except for pectin and mucin which were added at 1% (w/v). Growth was determined over a 48-h period using Tecan Infinite 50 (Tecan Ltd, UK) microplate spectrophotometer at OD_595_. Experiments were performed in biologically independent triplicates, and the plate reader measurements were taken automatically every 15 min following 60 s of shaking at normal speed. Due to the expected drop in initial OD values (i.e. recorded between T_0_ and T_1_) growth data were expressed as mean of the replicates between T_2_ (30 min) and T_end_ (48-h).

### High-performance anion-exchange chromatography (HPAEC)

Mono-, di- and oligo- saccharides present in the spent media samples were analyzed on a Dionex ICS-5000 HPAEC system operated by the Chromeleon software version 7 (Dionex, Thermo Scientific). Samples were bound to a Dionex CarboPac PA1 (Thermo Scientific) analytical column (2 × 250 mm) in combination with a CarboPac PA1 guard column (2 × 50 mm), equilibrated with 0.1 M NaOH. Carbohydrates were detected by pulsed amperometric detection (PAD). The system was run at a flow rate of 0.25 mL/min. The separation was done using a stepwise gradient going from 0.1 M NaOH to 0.1 M NaOH–0.1 M sodium acetate (NaOAc) over 10 min, 0.1 M NaOH–0.3 M NaOAc over 25 min followed by a 5 min exponential gradient to 1 M NaOAc, before reconditioning with 0.1 M NaOH for 10 min. Commercial glucose, cellobiose, fucose, lactose and lacto-*N*-neotetraose (LNnT) were used as external standards.

### Proteomics

*B. longum* subsp. *longum* strain 25 (B_25) was grown in triplicate in mMRS supplemented with cysteine HCl at 0.05% and 2% (w/v) glucose, cellobiose or LNnT as a sole carbon source. *B. longum* subsp. *longum* strain 71 (B_71) was grown in triplicate in mMRS supplemented with cysteine HCl at 0.05% and either 2% (w/v) glucose or 2’-fucosyllactose (2’-FL) as a sole carbon source. Cell pellets from 50 mL samples (at the mid-exponential growth phase) were collected by centrifugation (4500 × g, 10 min, 4 °C) and washed three times with PBS pH 7.4. Cells were resuspended in 50 mM Tris-HCl pH 8.4 and disrupted by bead-beating in three 60 s cycles using a FastPrep24 (MP Biomedicals, CA). Protein concentration was determined using a Bradford protein assay (Bio-Rad, Germany). Protein samples, containing 50 μg total protein, were separated by SDS-PAGE with a 10% Mini-PROTEAN gel (Bio-Rad Laboratories, CA) and then stained with Coomassie brilliant blue R250. The gel was cut into five slices, after which proteins were reduced, alkylated, and in-gel digested as previously described (*95*). Peptides were dissolved in 2% acetonitrile containing 0.1% trifluoroacetic acid and desalted using C18 ZipTips (Merck Millipore, Germany). Each sample was independently analysed on a Q-Exactive hybrid quadrupole-orbitrap mass spectrometer (Thermo Scientific) equipped with a nano-electrospray ion source. MS and MS/MS data were acquired using Xcalibur (v.2.2 SP1). Spectra were analysed using MaxQuant 1.6.1.0 (*96*) and searched against a sample-specific database generated from the B_25 and B_71 genomes. Proteins were quantified using the MaxLFQ algorithm (*97*). The enzyme specificity was set to consider tryptic peptides and two missed cleavages were allowed. Oxidation of methionine, N-terminal acetylation and deamidation of asparagine and glutamine and formation of pyro-glutamic acid at N-terminal glutamines were used as variable modifications, whereas carbamidomethylation of cysteine residues was used as a fixed modification. All identifications were filtered in order to achieve a protein false discovery rate (FDR) of 1% using the target-decoy strategy. A protein was considered confidently identified if it was detected in at least two of the three biological replicates in at least one glycan substrate. The MaxQuant output was further explored in Perseus v.1.6.1.1 (*98*). The proteomics data have been deposited to the ProteomeXchange Consortium (http://proteomecentral.proteomexchange.org) via the PRIDE partner repository with dataset identifier PXD017277.

## Acknowledgments

We would like to thank Glycom A/S for the kind donation of purified HMOs: 2′-FL and LNnT. The authors would also like to thank Prof Rob Kingsley and Mr Shabhonam Caim for technical support and advice. This work was funded by a Wellcome Trust Investigator Award (no. 100/974/C/13/Z); a BBSRC Norwich Research Park Bioscience Doctoral Training grant no. BB/M011216/1 (supervisor LJH, student MK); an Institute Strategic Programme Gut Microbes and Health grant no. BB/R012490/1 and its constituent projects BBS/E/F/000PR10353 and BBS/E/F/000PR10356; and an Institute Strategic Programme Gut Health and Food Safety grant no. BB/J004529/1 to LJH. LH was in receipt of a Medical Research Council Intermediate Research Fellowship in Data Science (UK MED-BIO, grant no. MR/L01632X/1). PBP and SLLR are grateful for support from The Research Council of Norway (FRIPRO program, PBP: 250479), as well as the European Research Commission Starting Grant Fellowship (awarded to PBP; 336355 - MicroDE). The funding bodies did not contribute to the design of the study, collection, analysis, and interpretation of data or in writing the manuscript.

## Author contributions

LJH, LH, ALM and MK designed the overall study. ALM provided the unique *B. longum* strain collection and extracted the DNA. MK prepared the DNA for WGS, performed all genomic analysis and visualisation, as well as growth studies. SLLR, PBP, LJH and MK planned metabolomics and proteomics studies. MK prepared samples for metabolomics and proteomics. SLLR and MK performed the metabolomics and proteomics experiments and SLLR analysed and visualised the resulting data. LJH and MK analysed the data, with input and discussion from LH, and drafted the manuscript. SLLR, PBP, LH and ALM provided providing further edits and co-writing of the final version. All authors read and approved the final manuscript.

## References

1. M. A. E. Lawson et al., Breast milk-derived human milk oligosaccharides promote Bifidobacterium interactions within a single ecosystem. ISME J 14, 635–648 (2020).

2. L. Wampach et al., Colonization and Succession within the Human Gut Microbiome by Archaea, Bacteria, and Microeukaryotes during the First Year of Life. Frontiers in microbiology 8, 738 (2017).

3. F. Backhed et al., Dynamics and Stabilization of the Human Gut Microbiome during the First Year of Life. Cell Host Microbe 17, 852 (2015).

4. M. G. de Aguero et al., The maternal microbiota drives early postnatal innate immune development. Science 351, 1296–1301 (2016).

5. A. Sivan et al., Commensal Bifidobacterium promotes antitumor immunity and facilitates anti-PD-L1 efficacy. Science 350, 1084–1089 (2015).

6. M. P. Heikkila, P. E. Saris, Inhibition of Staphylococcus aureus by the commensal bacteria of human milk. Journal of applied microbiology 95, 471–478 (2003).

7. M. Aaboud et al., Search for High-Mass Resonances Decaying to taunu in pp Collisions at sqrt[s]=13 TeV with the ATLAS Detector. Phys Rev Lett 120, 161802 (2018).

8. D. A. Sela et al., The genome sequence of Bifidobacterium longum subsp infantis reveals adaptations for milk utilization within the infant microbiome. P Natl Acad Sci USA 105, 18964–18969 (2008).

9. A. Marcobal, J. L. Sonnenburg, Human milk oligosaccharide consumption by intestinal microbiota. Clin Microbiol Infec 18, 12–15 (2012).

10. T. Thongaram, J. L. Hoeflinger, J. Chow, M. J. Miller, Human milk oligosaccharide consumption by probiotic and human-associated bifidobacteria and lactobacilli. Journal of dairy science 100, 7825–7833 (2017).

11. H. Renz, P. Brandtzaeg, M. Hornef, The impact of perinatal immune development on mucosal homeostasis and chronic inflammation. Nat Rev Immunol 12, 9–23 (2012).

12. P. J. Turnbaugh et al., An obesity-associated gut microbiome with increased capacity for energy harvest. Nature 444, 1027–1031 (2006).

13. T. Olszak et al., Microbial Exposure During Early Life Has Persistent Effects on Natural Killer T Cell Function. Science 336, 489–493 (2012).

14. N. A. Bokulich et al., Antibiotics, birth mode, and diet shape microbiome maturation during early life. Sci Transl Med 8, (2016).

15. W. H. W. Tang, T. Kitai, S. L. Hazen, Gut Microbiota in Cardiovascular Health and Disease. Circ Res 120, 1183–1196 (2017).

16. Q. Feng et al., Gut microbiome development along the colorectal adenoma-carcinoma sequence. Nat Commun 6, (2015).

17. Y. Shao et al., Stunted microbiota and opportunistic pathogen colonization in caesarean-section birth. Nature 574, 117–121 (2019).

18. A. O’Sullivan, M. Farver, J. T. Smilowitz, The Influence of Early Infant-Feeding Practices on the Intestinal Microbiome and Body Composition in Infants. Nutr Metab Insights 8, 1–9 (2015).

19. R. Martin et al., Early-Life Events, Including Mode of Delivery and Type of Feeding, Siblings and Gender, Shape the Developing Gut Microbiota. PLoS One 11, e0158498 (2016).

20. L. T. Stiemsma, K. B. Michels, The Role of the Microbiome in the Developmental Origins of Health and Disease. Pediatrics 141, (2018).

21. S. Ip et al., Breastfeeding and maternal and infant health outcomes in developed countries. Evid Rep Technol Assess (Full Rep), 1–186 (2007).

22. U. N. Das, Breastfeeding prevents type 2 diabetes mellitus: but, how and why? Am J Clin Nutr 85, 1436–1437 (2007).

23. J. A. Ortega-Garcia et al., Full Breastfeeding and Obesity in Children: A Prospective Study from Birth to 6 Years. Child Obes 14, 327–337 (2018).

24. J. D. Forbes et al., Association of Exposure to Formula in the Hospital and Subsequent Infant Feeding Practices With Gut Microbiota and Risk of Overweight in the First Year of Life. Jama Pediatr 172, (2018).

25. K. James, M. O. Motherway, F. Bottacini, D. van Sinderen, Bifidobacterium breve UCC2003 metabolises the human milk oligosaccharides lacto-N-tetraose and lacto-N-neo-tetraose through overlapping, yet distinct pathways. Sci Rep 6, 38560 (2016).

26. T. Katayama, Host-derived glycans serve as selected nutrients for the gut microbe: human milk oligosaccharides and bifidobacteria. Biosci Biotech Bioch 80, 621–632 (2016).

27. D. Garrido et al., A novel gene cluster allows preferential utilization of fucosylated milk oligosaccharides in Bifidobacterium longum subsp longum SC596. Sci Rep-Uk 6, (2016).

28. C. Milani et al., Bifidobacteria exhibit social behavior through carbohydrate resource sharing in the gut. Sci Rep-Uk 5, (2015).

29. J. E. Koenig et al., Succession of microbial consortia in the developing infant gut microbiome. Proc Natl Acad Sci U S A 108 Suppl 1, 4578–4585 (2011).

30. S. McKeen et al., Infant Complementary Feeding of Prebiotics for the Microbiome and Immunity. Nutrients 11, (2019).

31. M. B. Roberfroid, Inulin-type fructans: functional food ingredients. J Nutr 137, 2493S–2502S (2007).

32. W. F. Broekaert et al., Prebiotic and other health-related effects of cereal-derived arabinoxylans, arabinoxylan-oligosaccharides, and xylooligosaccharides. Crit Rev Food Sci Nutr 51, 178–194 (2011).

33. S. Hald et al., Effects of Arabinoxylan and Resistant Starch on Intestinal Microbiota and Short-Chain Fatty Acids in Subjects with Metabolic Syndrome: A Randomised Crossover Study. Plos One 11, (2016).

34. A. Riviere, M. Selak, D. Lantin, F. Leroy, L. De Vuyst, Bifidobacteria and Butyrate-Producing Colon Bacteria: Importance and Strategies for Their Stimulation in the Human Gut. Frontiers in microbiology 7, 979 (2016).

35. M. X. Maldonado-Gomez et al., Stable Engraftment of Bifidobacterium longum AH1206 in the Human Gut Depends on Individualized Features of the Resident Microbiome. Cell Host Microbe 20, 515–526 (2016).

36. K. Oki et al., Long-term colonization exceeding six years from early infancy of Bifidobacterium longum subsp. longum in human gut. Bmc Microbiol 18, (2018).

37. P. Mattarelli, C. Bonaparte, B. Pot, B. Biavati, Proposal to reclassify the three biotypes of Bifidobacterium longum as three subspecies: Bifidobacterium longum subsp. longum subsp. nov., Bifidobacterium longum subsp. infantis comb. nov. and Bifidobacterium longum subsp. suis comb. nov. Int J Syst Evol Microbiol 58, 767–772 (2008).

38. E. Yanokura et al., Subspeciation of Bifidobacterium longum by multilocus approaches and amplified fragment length polymorphism: Description of B. longum subsp. suillum subsp. nov., isolated from the faeces of piglets. Syst Appl Microbiol 38, 305–314 (2015).

39. F. Turroni et al., Exploring the Diversity of the Bifidobacterial Population in the Human Intestinal Tract. Appl Environ Microb 75, 1534–1545 (2009).

40. F. Turroni et al., Diversity of Bifidobacteria within the Infant Gut Microbiota. Plos One 7, (2012).

41. D. Garrido, D. Barile, D. A. Mills, A molecular basis for bifidobacterial enrichment in the infant gastrointestinal tract. Adv Nutr 3, 415S–421S (2012).

42. T. Odamaki et al., Genomic diversity and distribution of Bifidobacterium longum subsp longum across the human lifespan. Sci Rep-Uk 8, (2018).

43. L. C. Roger, A. L. McCartney, Longitudinal investigation of the faecal microbiota of healthy full-term infants using fluorescence in situ hybridization and denaturing gradient gel electrophoresis. Microbiol-Sgm 156, 3317–3328 (2010).

44. L. C. Roger, A. Costabile, D. T. Holland, L. Hoyles, A. L. McCartney, Examination of faecal Bifidobacterium populations in breast- and formula-fed infants during the first 18 months of life. Microbiol-Sgm 156, 3329–3341 (2010).

45. H. Ichinose, M. Yoshida, Z. Fujimoto, S. Kaneko, Characterization of a modular enzyme of exo-1,5-alpha-L-arabinofuranosidase and arabinan binding module from Streptomyces avermitilis NBRC14893. Appl Microbiol Biotechnol 80, 399–408 (2008).

46. S. Ahmed et al., A novel alpha-L-arabinofuranosidase of family 43 glycoside hydrolase (Ct43Araf) from Clostridium thermocellum. PLoS One 8, e73575 (2013).

47. H. Makino et al., Transmission of intestinal Bifidobacterium longum subsp. longum strains from mother to infant, determined by multilocus sequencing typing and amplified fragment length polymorphism. Appl Environ Microbiol 77, 6788–6793 (2011).

48. H. Makino et al., Mother-to-infant transmission of intestinal bifidobacterial strains has an impact on the early development of vaginally delivered infant’s microbiota. PLoS One 8, e78331 (2013).

49. C. Milani et al., Exploring Vertical Transmission of Bifidobacteria from Mother to Child. Appl Environ Microbiol 81, 7078–7087 (2015).

50. K. Pokusaeva, G. F. Fitzgerald, D. van Sinderen, Carbohydrate metabolism in Bifidobacteria. Genes Nutr 6, 285–306 (2011).

51. M. Ventura et al., Genome-scale analyses of health-promoting bacteria: probiogenomics. Nat Rev Microbiol 7, 61–71 (2009).

52. D. A. Sela, D. A. Mills, Nursing our microbiota: molecular linkages between bifidobacteria and milk oligosaccharides. Trends Microbiol 18, 298–307 (2010).

53. G. Davies, B. Henrissat, Structures and Mechanisms of Glycosyl Hydrolases. Structure 3, 853–859 (1995).

54. A. H. Viborg et al., Biochemical and kinetic characterisation of a novel xylooligosaccharide-upregulated GH43 beta-D-xylosidase/alpha-L-arabinofuranosidase (BXA43) from the probiotic Bifidobacterium animalis subsp lactis BB-12. Amb Express 3, (2013).

55. C. Milani et al., Genomics of the Genus Bifidobacterium Reveals Species-Specific Adaptation to the Glycan-Rich Gut Environment. Appl Environ Microb 82, 980–991 (2016).

56. S. K. Kang et al., Three forms of thermostable lactose-hydrolase from Thermus sp IB-21: cloning, expression, and enzyme characterization. J Biotechnol 116, 337–346 (2005).

57. S. W. A. Hinz, L. A. M. van den Broek, G. Beldman, J. P. Vincken, A. G. J. Voragen, Beta-galactosidase from Bifidobacterium adolescentis DSM20083 prefers beta(1,4)-galactosides over lactose. Appl Microbiol Biot 66, 276–284 (2004).

58. H. Suzuki, A. Murakami, K. Yoshida, Motif-guided identification of a glycoside hydrolase family 1 alpha-L-arabinofuranosidase in Bifidobacterium adolescentis. Biosci Biotechnol Biochem 77, 1709–1714 (2013).

59. V. Ambrogi et al., Characterization of GH2 and GH42 beta-galactosidases derived from bifidobacterial infant isolates. Amb Express 9, 9 (2019).

60. D. Garrido et al., Comparative transcriptomics reveals key differences in the response to milk oligosaccharides of infant gut-associated bifidobacteria. Sci Rep 5, 13517 (2015).

61. D. A. Sela et al., Bifidobacterium longum subsp. infantis ATCC 15697 alpha-fucosidases are active on fucosylated human milk oligosaccharides. Appl Environ Microbiol 78, 795–803 (2012).

62. M. Kitaoka, Bifidobacterial enzymes involved in the metabolism of human milk oligosaccharides. Adv Nutr 3, 422S–429S (2012).

63. S. W. Hinz, M. I. Pastink, L. A. van den Broek, J. P. Vincken, A. G. Voragen, Bifidobacterium longum endogalactanase liberates galactotriose from type I galactans. Appl Environ Microbiol 71, 5501–5510 (2005).

64. C. Yamada et al., Molecular Insight into Evolution of Symbiosis between Breast-Fed Infants and a Member of the Human Gut Microbiome Bifidobacterium longum. Cell Chem Biol 24, 515-+ (2017).

65. D. Watson et al., Selective carbohydrate utilization by lactobacilli and bifidobacteria. Journal of applied microbiology 114, 1132–1146 (2013).

66. S. Arboleya et al., Gene-trait matching across the Bifidobacterium longum pan-genome reveals considerable diversity in carbohydrate catabolism among human infant strains. Bmc Genomics 19, (2018).

67. S. Mills, C. Stanton, J. A. Lane, G. J. Smith, R. P. Ross, Precision Nutrition and the Microbiome, Part I: Current State of the Science. Nutrients 11, (2019).

68. J. M. Andersen et al., Transcriptional analysis of oligosaccharide utilization by Bifidobacterium lactis Bl-04. Bmc Genomics 14, 312 (2013).

69. S. Yan et al., Functional and structural characterization of a beta-glucosidase involved in saponin metabolism from intestinal bacteria. Biochem Biophys Res Commun 496, 1349–1356 (2018).

70. T. Pozzo, J. L. Pasten, E. N. Karlsson, D. T. Logan, Structural and functional analyses of beta-glucosidase 3B from Thermotoga neapolitana: a thermostable three-domain representative of glycoside hydrolase 3. J Mol Biol 397, 724–739 (2010).

71. R. N. Florindo et al., Structural and biochemical characterization of a GH3 beta-glucosidase from the probiotic bacteria Bifidobacterium adolescentis. Biochimie 148, 107–115 (2018).

72. E. Ozcan, D. A. Sela, Inefficient Metabolism of the Human Milk Oligosaccharides Lacto-N-tetraose and Lacto-N-neotetraose Shifts Bifidobacterium longum subsp. infantis Physiology. Front Nutr 5, (2018).

73. K. Mikami, M. Kimura, H. Takahashi, Influence of maternal bifidobacteria on the development of gut bifidobacteria in infants. Pharmaceuticals (Basel) 5, 629–642 (2012).

74. S. Lax et al., Longitudinal analysis of microbial interaction between humans and the indoor environment. Science 345, 1048–1052 (2014).

75. J. Dworkin, R. Losick, Linking nutritional status to gene activation and development. Genes Dev 15, 1051–1054 (2001).

76. J. Slager, J. W. Veening, Hard-Wired Control of Bacterial Processes by Chromosomal Gene Location. Trends Microbiol 24, 788–800 (2016).

77. D. E. Wood, S. L. Salzberg, Kraken: ultrafast metagenomic sequence classification using exact alignments. Genome Biol 15, (2014).

78. S. Chen, Y. Zhou, Y. Chen, J. Gu, fastp: an ultra-fast all-in-one FASTQ preprocessor. Bioinformatics 34, i884–i890 (2018).

79. A. Bankevich et al., SPAdes: a new genome assembly algorithm and its applications to single-cell sequencing. J Comput Biol 19, 455–477 (2012).

80. T. Seemann, Prokka: rapid prokaryotic genome annotation. Bioinformatics 30, 2068–2069 (2014).

81. L. Pritchard, R. H. Glover, S. Humphris, J. G. Elphinstone, I. K. Toth, Genomics and taxonomy in diagnostics for food security: soft-rotting enterobacterial plant pathogens. Anal Methods-Uk 8, 12–24 (2016).

82. J. Chun et al., Proposed minimal standards for the use of genome data for the taxonomy of prokaryotes. Int J Syst Evol Micr 68, 461–466 (2018).

83. A. J. Page et al., Roary: rapid large-scale prokaryote pan genome analysis. Bioinformatics 31, 3691–3693 (2015).

84. K. Katoh, J. Rozewicki, K. D. Yamada, MAFFT online service: multiple sequence alignment, interactive sequence choice and visualization. Brief Bioinform 20, 1160–1166 (2019).

85. T. Seemann. (2015).

86. N. J. Croucher et al., Rapid phylogenetic analysis of large samples of recombinant bacterial whole genome sequences using Gubbins. Nucleic Acids Res 43, (2015).

87. G. Talavera, J. Castresana, Improvement of phylogenies after removing divergent and ambiguously aligned blocks from protein sequence alignments. Systematic Biol 56, 564–577 (2007).

88. M. N. Price, P. S. Dehal, A. P. Arkin, FastTree 2-Approximately Maximum-Likelihood Trees for Large Alignments. Plos One 5, (2010).

89. T. Seemann, A. J. Page, F. Klotzl. (2017).

90. O. Brynildsrud, J. Bohlin, L. Scheffer, V. Eldholm, Rapid scoring of genes in microbial pan-genome-wide association studies with Scoary. Genome Biol 17, 238 (2016).

91. J. Huerta-Cepas et al., Fast Genome-Wide Functional Annotation through Orthology Assignment by eggNOG-Mapper. Mol Biol Evol 34, 2115–2122 (2017).

92. H. Zhang et al., dbCAN2: a meta server for automated carbohydrate-active enzyme annotation. Nucleic Acids Res 46, W95–W101 (2018).

93. M. Csuros, I. Miklos, A probabilistic model for gene content evolution with duplication, loss, and horizontal transfer. Lect Notes Comput Sc 3909, 206–220 (2006).

94. A. J. Page et al., SNP-sites: rapid efficient extraction of SNPs from multi-FASTA alignments. Microb Genomics 2, (2016).

95. M. O. Arntzen, I. L. Karlskas, M. Skaugen, V. G. H. Eijsink, G. Mathiesen, Proteomic Investigation of the Response of Enterococcus faecalis V583 when Cultivated in Urine. Plos One 10, (2015).

96. J. Cox, M. Mann, MaxQuant enables high peptide identification rates, individualized p.p.b.-range mass accuracies and proteome-wide protein quantification. Nat Biotechnol 26, 1367–1372 (2008).

97. J. Cox et al., Accurate Proteome-wide Label-free Quantification by Delayed Normalization and Maximal Peptide Ratio Extraction, Termed MaxLFQ. Mol Cell Proteomics 13, 2513–2526 (2014).

98. S. Tyanova et al., The Perseus computational platform for comprehensive analysis of (prote)omics data. Nat Methods 13, 731–740 (2016).

