## Supplementary figures and images for "Succession of *Bifidobacterium longum* strains in response to the changing early-life nutritional environment reveals specific adaptations to distinct dietary substrates"

### Supplementary Figure 1

#### Slide 1
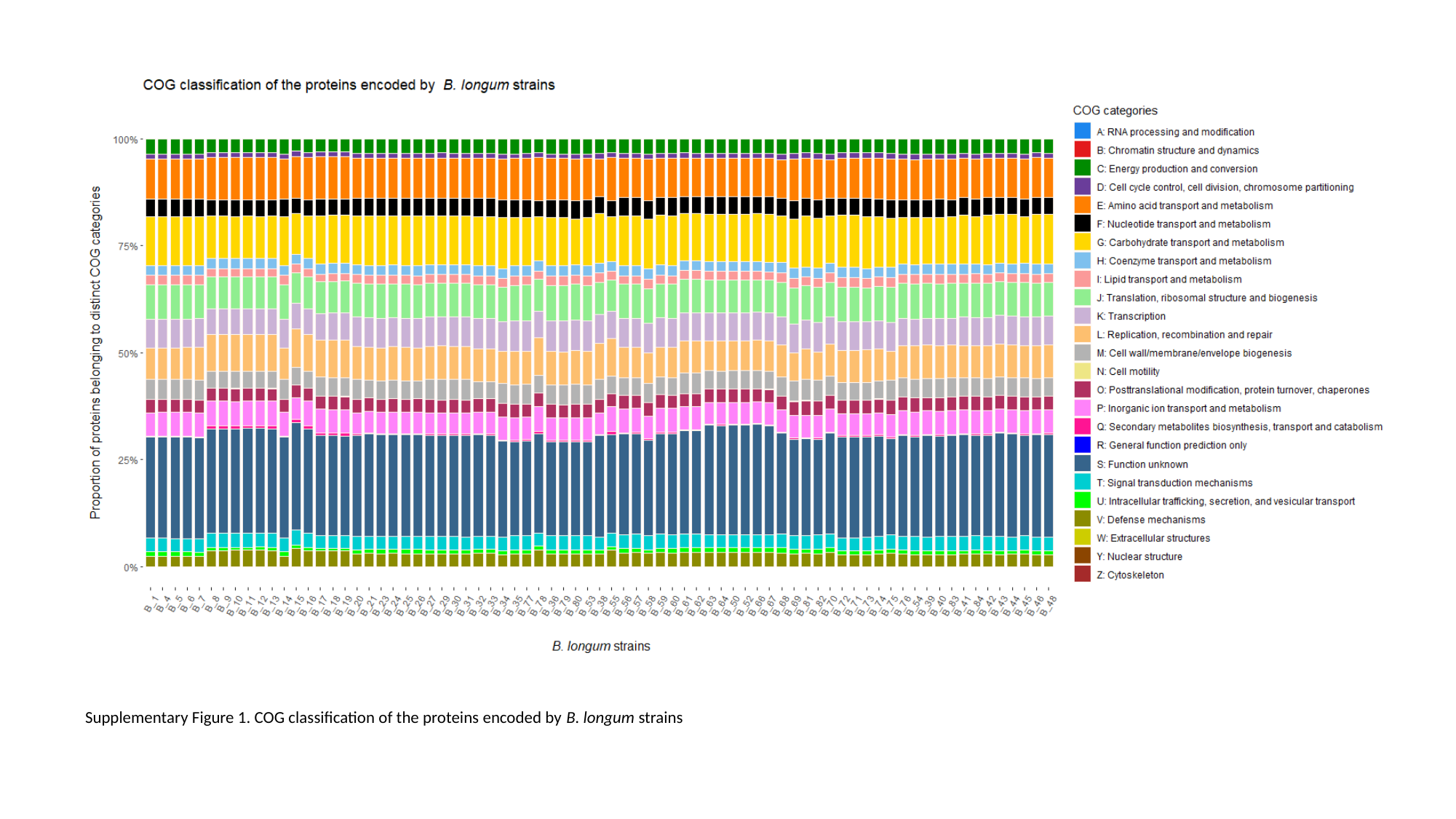

Supplementary Figure 1. COG classification of the proteins encoded by B. longum strains
